# Role of cation-chloride cotransporters, Na/K-pump, and channels in cell water/ionic balance regulation under hyperosmolar conditions: in silico and experimental studies of opposite RVI and AVD responses of U937 cells to hyperosmolar media

**DOI:** 10.1101/2021.11.15.468591

**Authors:** Valentina E. Yurinskaya, Alexey A. Vereninov

## Abstract

The work provides a modern mathematical description of animal cell electrochemical system under a balanced state and during the transition caused by an increase in external osmolarity, considering all the main ionic pathways in the cell membrane: the sodium pump, K^+^, Na^+^, Cl^-^ electroconductive channels and cotransporters NC, KC, and NKCC. The description is applied to experimental data obtained on U937 cells cultured in suspension, which allows the required assays to be performed, including determination of cell water content using buoyant density, cell ion content using flame photometry, and optical methods using flow cytometry. The study of these cells can serve as a useful model for understanding the general mechanisms of regulation of cellular water and ionic balance, which cannot be properly analyzed in many important practical cases, such as ischemic disturbance of cellular ionic and water balance, when cells cannot be isolated. An essential part of the results is the developed software supplied with an executable file, which allows researchers with no programming experience to calculate unidirectional fluxes of monovalent ions through separate pathways and ion-electrochemical gradients that move ions through them, which is important for studying the functional expression of channels and transporters. It is shown how the developed approach is used to reveal changes in channels and transporters underlying the RVI and AVD responses to the hyperosmolar medium in the studied living U937 cells.

## Introduction

Many processes at the physiological, proteomic, and genomic levels are triggered in cells already in the first hour after the increase in the osmolarity of the external environment, which is often called “osmotic stress” (Burg et al., 2007; Lambert et al., 2008; Hoffmann et al., 2009; Koivusalo et al., 2009; Wang et al., 2014). Rapid osmotic shrinkage of cells is usually accompanied by a regulatory increase in volume, RVI, and, with some delay, an oppositely directed decrease in volume, AVD, which is associated with the initiation of apoptosis (Yurinskaya et al., 2012). Monovalent ions redistribution mechanisms which cause RVI and AVD remain insufficiently studied. There is no quantitative description of transient processes in the cell electrochemical system caused by replacing the isoosmolar medium with a hyperosmolar medium, which would consider, in addition to the sodium pump and electrically conductive channels, all the main types of cation-chloride cotransporters. We tried to fill this gap using a mathematical analysis of the complex interdependence of ion fluxes via the main pathways across the cell membrane and an experimental study of living U937 cells included determination of cell water content by buoyant density, cell ion content using flame photometry, and optical methods using flow cytometry. It is found by this way that in U937 cells studied as example: (1) an effect like RVI can take place in hyperosmolar media with addition of NaCl without changes in the membrane channels and transporters if certain cation-chloride cotransporters present in the cell membrane, (2) time-dependent decrease in cell volume, such as AVD, can occur in the hyperosmolar medium of sucrose without changes in membrane ion channels and transporters; (3) the response of living cells to a hypoosmolar challenge is more complex than the response of their electrochemical model due to regulation of transporters by intracellular signaling mechanisms and due to changes in the content of intracellular impermeable osmolytes. These specific effects are identified by eliminating the “physical” effects found by mathematical analysis of the entire electrochemical system of the cell.

## Materials and methods

### Reagents

RPMI 1640 medium and fetal bovine serum (FBS, HyClone Standard) were purchased from Biolot (Russia). Ouabain was from Sigma-Aldrich (Germany), Percoll was purchased from Pharmacia (Sweden). The isotope ^36^Cl^−^ was from “Isotope” (Russia). Salts and sucrose were of analytical grade and were from Reachem (Russia).

### Cell cultures and solutions

Lymphoid cell lines U937, K562, and Jurkat from the Russian Cell Culture Collection (Institute of Cytology, Russian Academy of Sciences) were studied. The cells were cultured in RPMI 1640 medium supplemented with 10% FBS at 37 °C and 5% CO_2_ and subcultured every 2-3 days. Cells with a culture density of approximately 1×10^6^ cells per ml were transferred to hyperosmolar rmedium with addition of 100 mM NaCl or 150-300 mM sucrose for 0-4 h. A stock solution of 1 mM NaCl or 2 mM sucrose in PBS was used to add to the standard RPMI medium to prepare a hyperosmolar medium. The osmolarity of solutions was checked with the Micro-osmometer Model 3320 (Advanced Instruments, USA). All the incubations were done at 37 °C.

### Determination of cell water and ion content

Details of the experimental methods used were described in our previous study (Yurinskaya et al., 2019-2021). Briefly, cell water content was estimated by the buoyant density of the cells in continuous Percoll gradient, intracellular K^+^, Na^+^ and Rb^+^ content was determined by flame emission on a Perkin-Elmer AA 306 spectrophotometer, the intracellular Cl^−^ was measured using a radiotracer ^36^Cl. The cell water content was calculated in ml per gram of protein as v_prot_ = (1 − ρ/ρ_dry_)/[0.72(ρ − 1)], where ρ is the measured buoyant density of the cells and ρ_dry_ is the density of the cell dry mass, the latter taken as 1.38 g/ml. The ratio of protein to dry mass was taken as 0.72. The cellular ion content was calculated in micromoles per gram of protein

### Statistical analysis

Experimental data are presented as the mean ± SEM. P < 0.05 (Student’s t test) was considered statistically significant. Statistical analysis for calculated data is not applicable.

### The mathematical background of the modeling

The mathematical model of the movement of monovalent ions across the cell membrane was like that used by Jakobsson (1980), and Lew with colleagues (Lew, Bookchin, 1986; Lew et al. 1991; Lew, 2000), as well as in our previous works (Vereninov et al., 2014, 2016; Yurinskaya et al., 2019, 2020, 2021). It accounts for the Na/K pump, electroconductive channels, cotransporters NC, KC, and NKCC. NKCC indicates the known cotransporters of the SLC12 family carrying monovalent ions with stoichiometry 1Na^+^:1K^+^:2Cl^−^ and KC and NC stand for cotransporters with stoichiometry 1K^+^:1Cl^−^ or 1Na^+^:1Cl^−^. The latter can be represented by a single protein, the thiazide-sensitive Na-Cl cotransporter (SLC12 family), or by coordinated operation of the exchangers Na/H, SLC9, and Cl/HCO_3_, SLC26 (Garcia-Soto and Grinstein, 1990). In the considered approach, the entire set of ion transport systems is replaced by a reduced number of ion pathways, determined thermodynamically, but not by their molecular structure. All the major pathways are subdivided into five subtypes by ion-driving force: ion channels, where the driving force is the transmembrane electrochemical potential difference for a single ion species; NKCC, NC, and KC cotransporters, where the driving force is the sum of the electrochemical potential differences for all partners; and the Na/K ATPase pump, where ion movement against electrochemical gradient is energized by ATP hydrolysis. This makes it possible to characterize the intrinsic properties of each pathway using a single rate coefficient. The using the model with single parameters for characterization of each ion pathways is quite sufficient for successful description of the homeostasis in real cells at a real accuracy of the current experimental data.

The basic equations are presented below. Symbols and definitions used are shown in **Table 1.** Two mandatory conditions of macroscopic electroneutrality and osmotic balance:

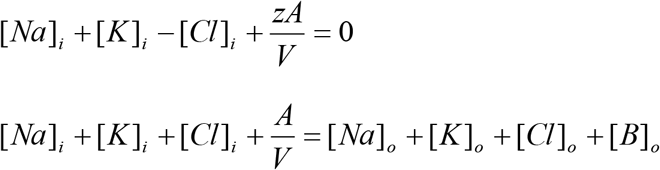

**Table 1.**
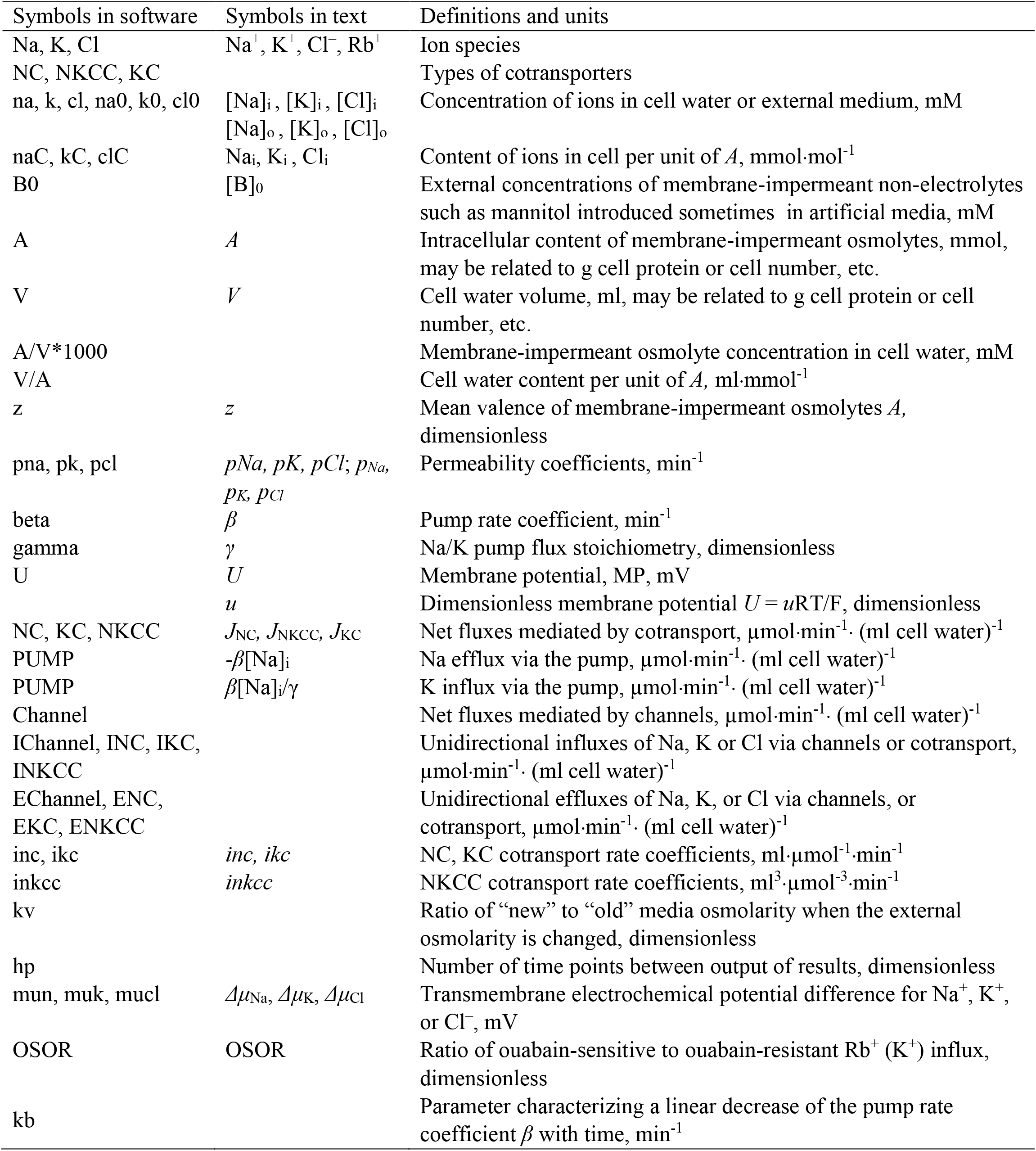
Symbols and definitions.

| Symbols in software | Symbols in text | Definitions and units |
| --- | --- | --- |
| Na, K, Cl | $\text{Na}^+$, $\text{K}^+$, $\text{Cl}^-$, $\text{Rb}^+$ | Ion species |
| NC, NKCC, KC |  | Types of cotransporters |
| na, k, cl, na0, k0, cl0 | $[\text{Na}]_i$, $[\text{K}]_i$, $[\text{Cl}]_i$<br>$[\text{Na}]_o$, $[\text{K}]_o$, $[\text{Cl}]_o$ | Concentration of ions in cell water or external medium, mM |
| naC, kC, clC | $\text{Na}_i$, $\text{K}_i$, $\text{Cl}_i$ | Content of ions in cell per unit of A, $\text{mmol}\cdot\text{mol}^{-1}$ |
| B0 | $[\text{B}]_0$ | External concentrations of membrane-impermeant non-electrolytes such as mannitol introduced sometimes in artificial media, mM |
| A | A | Intracellular content of membrane-impermeant osmolytes, mmol, may be related to g cell protein or cell number, etc. |
| V | V | Cell water volume, ml, may be related to g cell protein or cell number, etc. |
| A/V*1000 |  | Membrane-impermeant osmolyte concentration in cell water, mM |
| V/A | | Cell water content per unit of A, $\text{ml}\cdot\text{mmol}^{-1}$ |
| z | z | Mean valence of membrane-impermeant osmolytes A, dimensionless |
| pna, pk, pcl | $p\text{Na}$, $p\text{K}$, $p\text{Cl}$; $p_{\text{Na}}$,<br>$p_{\text{K}}$, $p_{\text{Cl}}$ | Permeability coefficients, $\text{min}^{-1}$ |
| beta | $\beta$ | Pump rate coefficient, $\text{min}^{-1}$ |
| gamma | $\gamma$ | Na/K pump flux stoichiometry, dimensionless |
| U | U | Membrane potential, MP, mV |
| | u | Dimensionless membrane potential $U = uRT/F$, dimensionless |
| NC, KC, NKCC | $J_{\text{NC}}$, $J_{\text{NKCC}}$, $J_{\text{KC}}$ | Net fluxes mediated by cotransport, $\mu\text{mol}\cdot\text{min}^{-1}\cdot(\text{ml cell water})^{-1}$ |
| PUMP | $-\beta[\text{Na}]_i$ | Na efflux via the pump, $\mu\text{mol}\cdot\text{min}^{-1}\cdot(\text{ml cell water})^{-1}$ |
| PUMP | $\beta[\text{Na}]_i/\gamma$ | K influx via the pump, $\mu\text{mol}\cdot\text{min}^{-1}\cdot(\text{ml cell water})^{-1}$ |
| Channel | | Net fluxes mediated by channels, $\mu\text{mol}\cdot\text{min}^{-1}\cdot(\text{ml cell water})^{-1}$ |
| IChannel, INC, IKC, INKCC | | Unidirectional influxes of Na, K or Cl via channels or cotransport, $\mu\text{mol}\cdot\text{min}^{-1}\cdot(\text{ml cell water})^{-1}$ |
| EChannel, ENC, EKC, ENKCC | | Unidirectional effluxes of Na, K, or Cl via channels, or cotransport, $\mu\text{mol}\cdot\text{min}^{-1}\cdot(\text{ml cell water})^{-1}$ |
| inc, ikc | $inc$, $ikc$ | NC, KC cotransport rate coefficients, $\text{ml}\cdot\mu\text{mol}^{-1}\cdot\text{min}^{-1}$ |
| inkcc | $inkcc$ | NKCC cotransport rate coefficients, $\text{ml}^3\cdot\mu\text{mol}^{-3}\cdot\text{min}^{-1}$ |
| kv |  | Ratio of “new” to “old” media osmolarity when the external osmolarity is changed, dimensionless |
| hp |  | Number of time points between output of results, dimensionless |
| mun, muk, mucl | $\Delta\mu_{\text{Na}}$, $\Delta\mu_{\text{K}}$, $\Delta\mu_{\text{Cl}}$ | Transmembrane electrochemical potential difference for $\text{Na}^+$, $\text{K}^+$, or $\text{Cl}^-$, mV |
| OSOR | OSOR | Ratio of ouabain-sensitive to ouabain-resistant $\text{Rb}^+$ ($\text{K}^+$) influx, dimensionless |
| kb | | Parameter characterizing a linear decrease of the pump rate coefficient $\beta$ with time, $\text{min}^{-1}$ |

The flux equations:

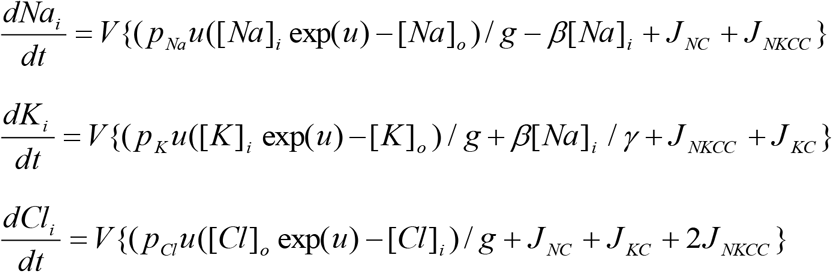

The left-hand sides of these three equations represent the rates of change of cell ion content. The right-hand sides express fluxes, where *u* is the dimensionless membrane potential related to the absolute values of membrane potential *U* (mV), as *U* = *u*RT/F = 26.7*u* for 37 °C and *g* = 1 − exp(*u*). The rate coefficients *p_Na_, p_K_, p_Cl_* characterizing channel ion transfer are similar to the Goldman’s coefficients. Fluxes *J_NC_, J_KC_, J_NKCC_* depend on internal and external ion concentrations as

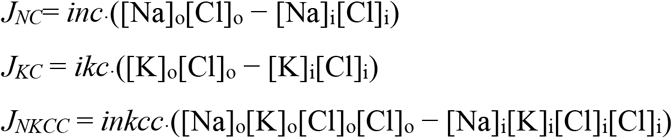

Here *inc, ikc*, and *inkcc* are the rate coefficients for cotransporters.

The rate coefficient of the sodium pump (*beta*) was calculated as the ratio of the Na^+^ pump efflux to the cell Na^+^ content where the Na^+^ pump efflux was estimated by ouabain-sensitive (OS) K^+^(Rb^+^) influx assuming proportions of [Rb]_o_ and [K]_o_, respectively, and Na/K pump flux stoichiometry of 3:2.

Transmembrane electrochemical potential differences for Na^+^, K^+^, and Cl^−^ were calculated as: *Δμ*_Na_ =26.7·ln([Na]_i_ /[Na]_o_)+*U*, *Δμ*_K_ =26.7·ln([K]_i_ /[K]_o_)+*U*, and *Δμ*_Cl_ =26.7·ln([Cl]_i_ /[Cl]_o_)-*U*, respectively. The algorithm of the numerical solution of the system of these equations is considered in detail in (Vereninov et al., 2014), the using of the executable file is illustrated more in (Yurinskaya et al., 2019). The problems in determination of the multiple parameters in a system with multiple variables like cell ionic homeostasis are discussed in more detail in (Yurinskaya et al., 2019, 2020). Some readers of our previous publications have expressed doubt that using our tool it is possible to obtain a unique set of parameters that provide an agreement between experimental and calculated data. Our mathematical comments on this matter can be found in Yurinskaya et al., 2019 (Notes Added in Response to SomeReaders).

The executable file of the BEZ02BC program used in this study with two auxiliary files is presented in the Supplement. It differs slightly from our previous executable file BEZ01B by replacing in the output table the columns prn, prk, prcl (time derivatives of concentrations) with the columns naC, kC, and clC representing intracellular content of Na^+^, K^+^, and Cl^−^.

## Results

### 1. Rearrangement of cell ionic homeostasis in hyperosmolar media, calculated for a system with different, but invariable in time, membrane parameters

Earlier it was shown that the same balanced intracellular concentrations of Na^+^, K^+^, Cl^-^, the content of cell water, and the coefficient of the pumping rate, as in the experiment, can be obtained in model with several sets of cotransporters (Yurinskaya and Vereninov, 2021). Despite the difficulties with increasing the number of parameters, the use of additional data on the action of specific inhibitors allows one to determine the required parameters (Yurinskaya et al., 2019, 2020). It was found also that the living U937 cells of the same established line can differ in real physiological experiment by the functional expression of cotransporters. For these reasons, in studying cell response to an increase in external osmolarity several sets of cotransporters are considered some of which have analogs among the living cells. The values of the permeability coefficients of the Na^+^, K^+^, and Cl^-^ channels corresponding to the balanced distribution of ions inevitably differ depending on the chosen cotransporters. The electrical, electrochemical potential differences and their derivatives also significantly differ depending on the specified cotransporters (**Table 2).** This is an example of when cotransporters with electrically neutral ion transport significantly affect the membrane potential in the system due to changes in the intracellular ion concentration (see difference between “voltagenic” and “amperogenic” ion transfers in Stanton, 1983).

**Table 2.** Basic characteristics of ion distribution, measured in living U937 cells equilibrated with a normal isotonic medium RPMI, and computed for models with different sets of parameters.

| Measured characteristics in normal RPMI medium |  |  |  |  |
| --- | --- | --- | --- | --- |
| [K]<sub>i</sub> 147, [Na]<sub>i</sub> 38, [Cl]<sub>i</sub> 45, A<sup>z</sup> 80 mM, |  |  |  |  |
| V/A 12.5 ml/mmol, Beta 0.039 min<sup>-1</sup>, OSOR 3.89, z -1.75 |  |  |  |  |
| Cotransporters assigned for balanced state in normal medium |  |  |  |  |
|  | NC | NC+KC | NC+NKCC | NC+KC+NKCC |
| *inc* | 3E-5 | 4.87E-5 | 3E-5 | 7E-5 |
| *ikc* | – | 6E-5 | – | 8E-5 |
| *inkcc* | – | – | 7E-9 | 8E-9 |
| Parameters computed for balanced state in normal medium |  |  |  |  |
| *pna* | 0.00382 | 0.00263 | 0.0043 | 0.0017 |
| *pk* | 0.02200 | 0.0165 | 0.0175 | 0.0115 |
| *pcl* | 0.00910 | 0.006 | 0.0139 | 0.011 |
| Computed characteristics in normal medium, mV |  |  |  |  |
| *U* | -44.7 | -49.3 | -37.6 | -45.0 |
| *mucl* | +19.4 | +24.0 | +12.3 | +19.8 |
| *mun* | -79.5 | -84.1 | -72.4 | -79.9 |
| *muk* | +41.6 | +37.0 | +48.7 | +41.3 |

Data related to the sample of the living cells U937 described in our 2021 article as Cells B are used as an example in this study.

**Figure 1** shows calculated transition to hyperosmolar media in cells with the identical initial intracellular Na^+^, K^+^, Cl^-^ concentrations (38, 147, 45 mM, respectively), cell water content, and the same sodium pump rate coefficient (0.039 min^-1^) but with different set of the assigned cotransporters. Computation shows the new ionic homeostasis is established with time both in sucrose (**Figure 1A-D**) and in NaCl hyperosmolar media (**Figure 1G-J)**. Mathematically, this follows from the feedback between the intracellular concentration of Na^+^ and the efflux of Na^+^ through the pump. The higher the intracellular Na^+^ concentration, the greater the outflow of the pump with the same pumping rate *beta*, that is, with the same intrinsic pump properties. As soon as a new water balance is established, the volume of cells in a hyperosmolar medium with additional 180 mM sucrose or 100 mM NaCl should be 0.63 (310/490) or 0.61 (310/510) of the volume in a medium with a normal osmolarity 310 mOsm. This follows simply from the equation of osmotic balance, which is achieved much faster than the ionic balance.

**Figure 1.**
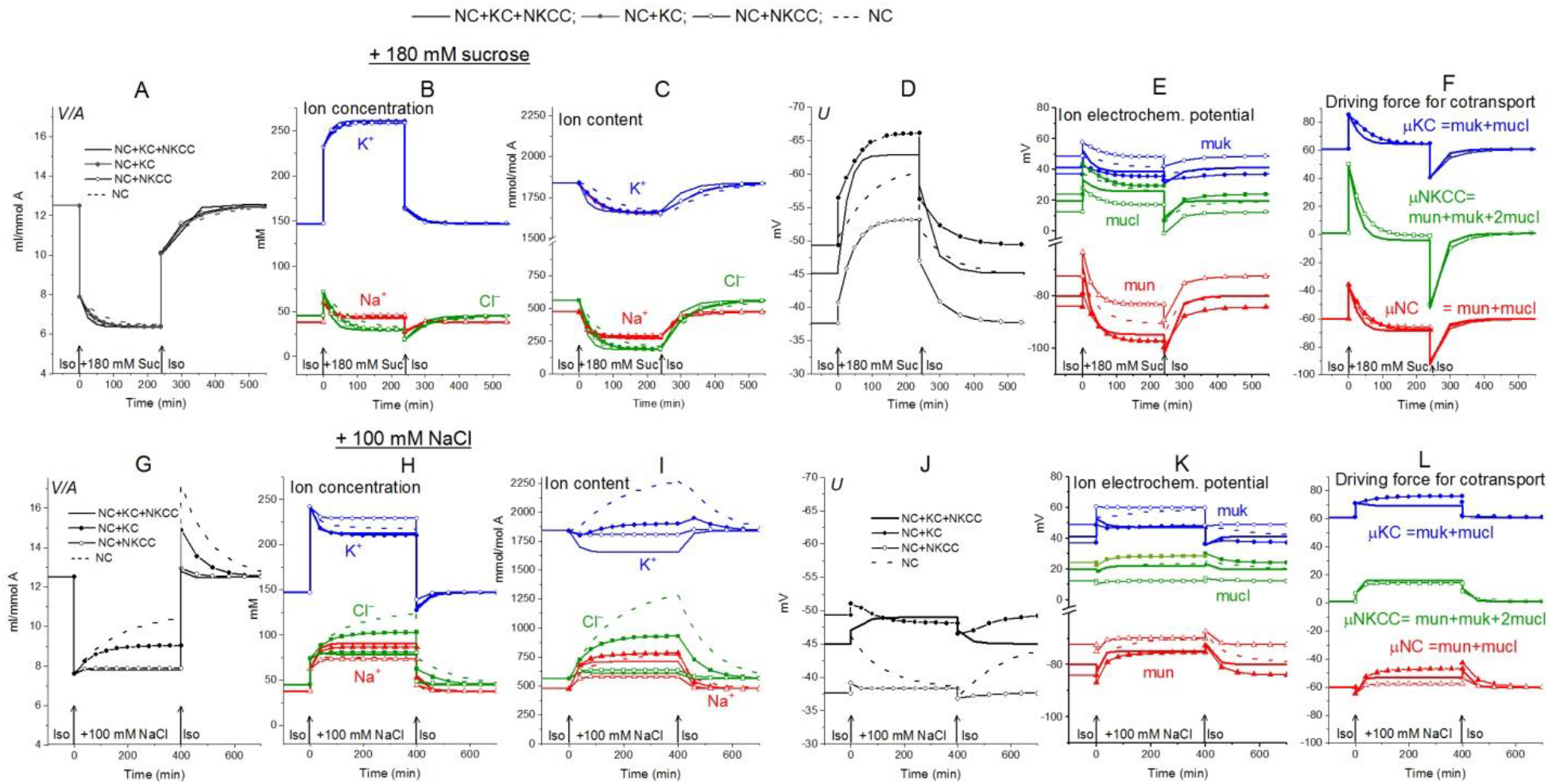
Rearrangement of ionic homeostasis following an increase in external osmolarity due to addition of 180 mM sucrose (**A-F**) or 100 mM NaCl (**G-L**) and a reverse transition to the normal medium, calculated for a model of U937 cells with different cotransporters and invariable in time parameters of channels and transporters like in U937 cells, equilibrated with the standard RPMI medium.

Computation shows how the transition to new cell water and ionic homeostasis can differ in NaCl and sucrose hyperosmolar media, and how this difference can depend on the cotransporters in the cell membrane (**Figure 1A, G, Table 3**). A time-dependent increase in cell volume, like RVI in living cells, occurs in a hyperosmolar medium supplemented with NaCl under appropriate conditions (V/V_initial_>1, marked in yellow in Table 3), while, in contrast, a decrease in cell volume increase, AVD (RVD), occurs in sucrose hyperosmolar medium (V/V_initial_<1, marked in blue in Table 3). Changes in cell volume are associated in all cases with changes in the K^+^, Na^+^ and Cl^-^ content of the corresponding sign. A new balanced state in the hyperosmolar medium with sucrose is achieved due to the approximately equal exit of K^+^ and Na^+^ from the cell. In the medium with 100 mM NaCl, changes in K^+^ and Na^+^, which underlie RVI, differ significantly depending on the set cotransporters. In the NC model, K^+^ uptake plays a major role in RVI, while in the NC+KC model, Na^+^ uptake dominates. When NKCC is present in the membrane, RVI is associated with Na^+^ uptake and small K^+^ release (**Table 3**). It is essential that in all cases the RVI-like effect in the hyperosmolar NaCl medium and the AVD-like effect in the hyperosmolar sucrose medium do not require any “regulatory” changes in the parameters of membrane channels and transporters. It is noteworthy that a significant RVI effect in the hyperosmolar NaCl medium is observed only in the system without the NKCC cotransporter (**Table 3, Figure 1G**). The difference in behavior of the cell ionic system in the sucrose and NaCl hyperosmolar solutions occurs because we are dealing with charged but not with electrically neutral osmolytes. Consideration of the model with the unchanged in time cell membrane parameters shows that the changes in the electrochemical potential difference moving K^+^, Na^+^ and Cl^-^ via corresponding pathways can be a solely determinant of the kinetics of the ion homeostasis rearrangement. The electrochemical potential differences for each of the ions, K^+^, Na^+^ and Cl^-^, in cells placed into sucrose and into NaCl hypertonic media can change by different way and even in the opposite directions as well as the magnitudes and signs of the forces driving ions via the NC, KC and NKCC cotransporters (**Figure 1E, F, K, L)**.

**Table 3.**
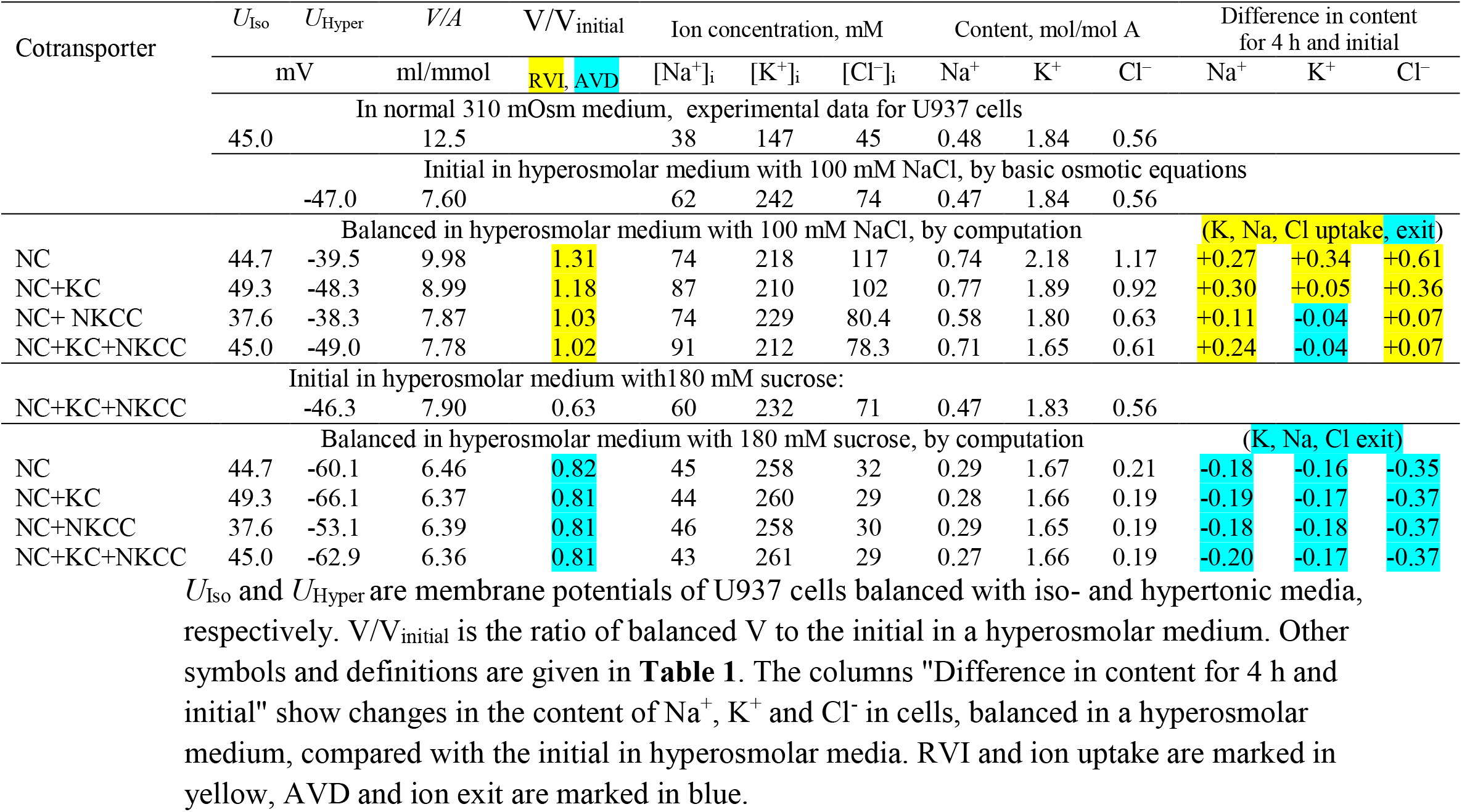
Changes in ionic homeostasis under new balanced state in hyperosmolar media with 100 mM NaCl or 180 mM sucrose, calculated for the U937 cell model with different sets of cotransporters and parameters corresponding to cells balanced with normal 310 mOsm medium.

The direction of the force driving ions through the pathways of cotransporters determines their role in the regulation of the ionic and water balance of the cell. Changes in the rate coefficients *inc* and *ikc* have approximately the same effect as changes in *pk, pna*, and *pcl*, that is, they zero out the electrochemical gradients of monovalent ions generated by the sodium pump and decrease the membrane potential. They are antagonists of the pump (Vereninov et al., 2014; Dmitriev et al., 2021). This is not the case in an increase in *inkcc* rate coefficient which shifts the system to the ion distribution when the driving force for the transport of ions through the NKCC is close to zero. In cells such as U937, as well as in many other cells, when they are balanced with the normal medium, the relationship between concentration ratio for K^+^, Na^+^ and Cl^-^ on the membrane correspond to zero driving force for the transport of ions through the NKCC (**Figire 1**). In this case NKCC cotransporter stabilizes the normal state of a cell. Variation in the rate coefficient *inkcc* has no effect on the state of the electrochemical system of a normal cell. The deviation of ion distribution from the standard generates non-zero μNKCC which declines with time to zero in the sucrose hyperosmolar medium but not in the NaCl hyperosmolar medium (**Figure 1**). An increase in μNKCC gradient generates the net fluxes of Na^+^, K^+^, and Cl^-^ via NKCC pathway. It looks like “activation” of NKCC transporter although the intrinsic properties of cotransporter, i.e., rate coefficient in the model is not changed. The increase in the net fluxes of K^+^, Na^+^ and Cl^-^ via NKCC caused by an increase in the μNKCC gradient is clearly seen if the dynamic of the unidirectional and net fluxes is considered which is shown in **Table 4.** The considered examples show that specific NKCC blockers like bumetanide and its analogues can have no effect on the entire ion homeostasis in normal cells. However, it would be mistaken to say that the NKCC is absent or “not-activated” and “silent”. The unidirectional fluxes via NKCC pathway can be significant and presence of this cotransporter can be revealed for example by measurement of the bumetanide-inhibitable Rb^+^ influx. The situation changes when a normal medium is replaced with the hyperosmolar one, especially with the NaCl hyperosmolar medium. The impact of the net fluxes of Na^+^, K^+^ and Cl^-^ via NKCC in the total flux balance increases significantly and inhibitors become effective.

**Table 4.**
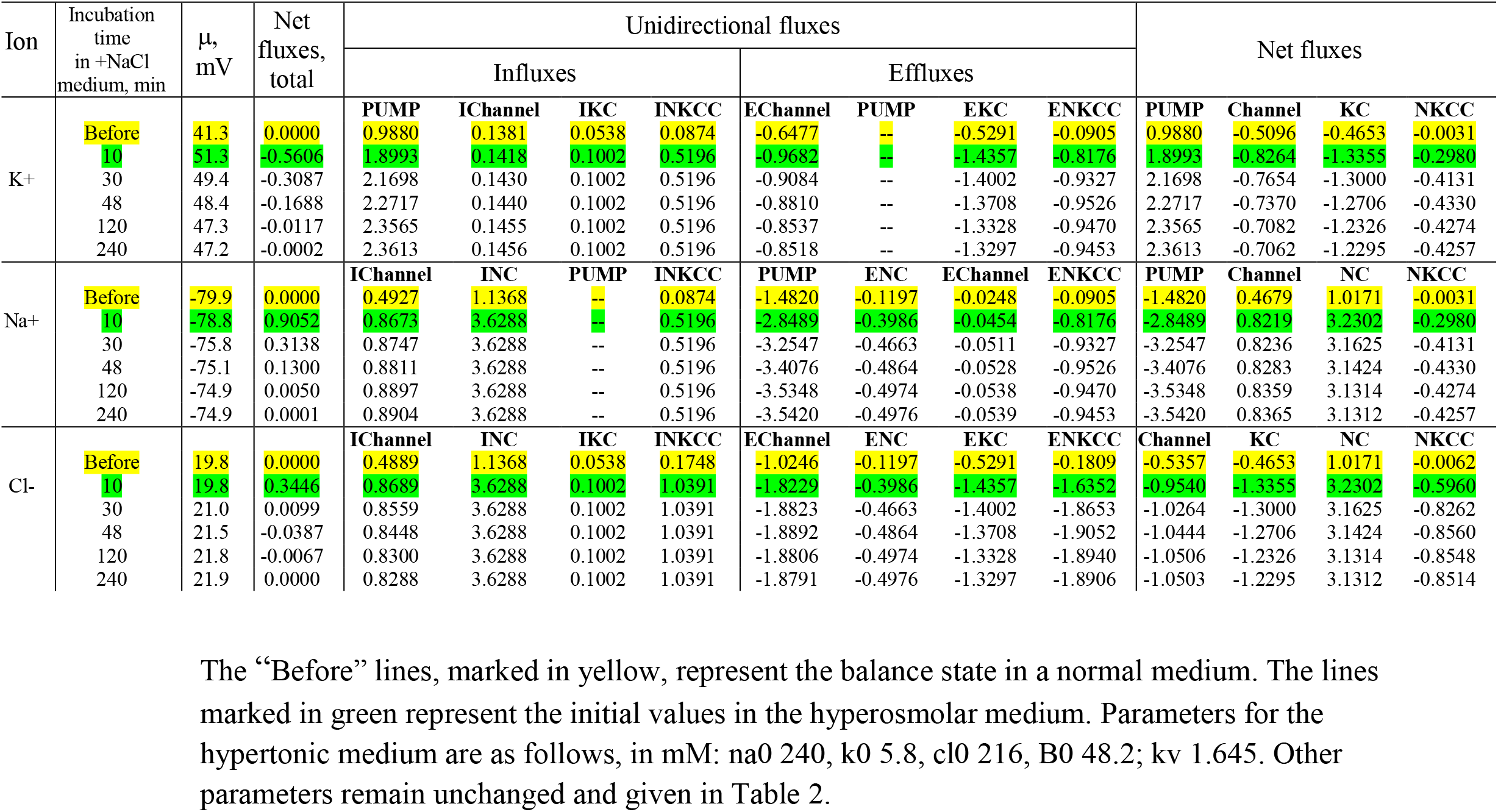
Dynamics of the net and unidirectional K^+^, Na^+^, and Cl^−^ fluxes in U937 cells during transition Iso-Hyper (+100 NaCl) calculated for the modelwith all main cotransporters and parameters like in cells U937 equilibrated with standard 310 mOsm medium.

Alteration of ionic homeostasis in hyperosmolar media is fully restored after the return of cells into the medium of normal osmolarity. (**Figures 1 and 2**). The kinetics of the Iso-Hyper-Iso transition, both direct and reverse, significantly depends on the set of cotransporters in the case of a NaCl hyperosmolar solution. This occurs mostly because of the different basic pCl at different cotransporters. At low pCl, the new balance is reached more slowly. We observed a similar effect when modeling changes in ionic homeostasis due to blockage of the sodium pump (Yurinskaya et al., 2019). The kinetics of the reverse Hyper-Iso transition for both sucrose and NaCl cases is at first glance similar and copies that of the direct Iso-Hyper transition. However, a more detailed analysis shows that there are certain differences between changes in the direct and reverse direction, and between the cases of sucrose and NaCl hyperosmolar solutions (**Figure 2**). As mentioned above, NKCC cotransport attenuates changes in homeostasis both in the hyperosmolar medium with sucrose and with additional NaCl.

**Figure 2.**
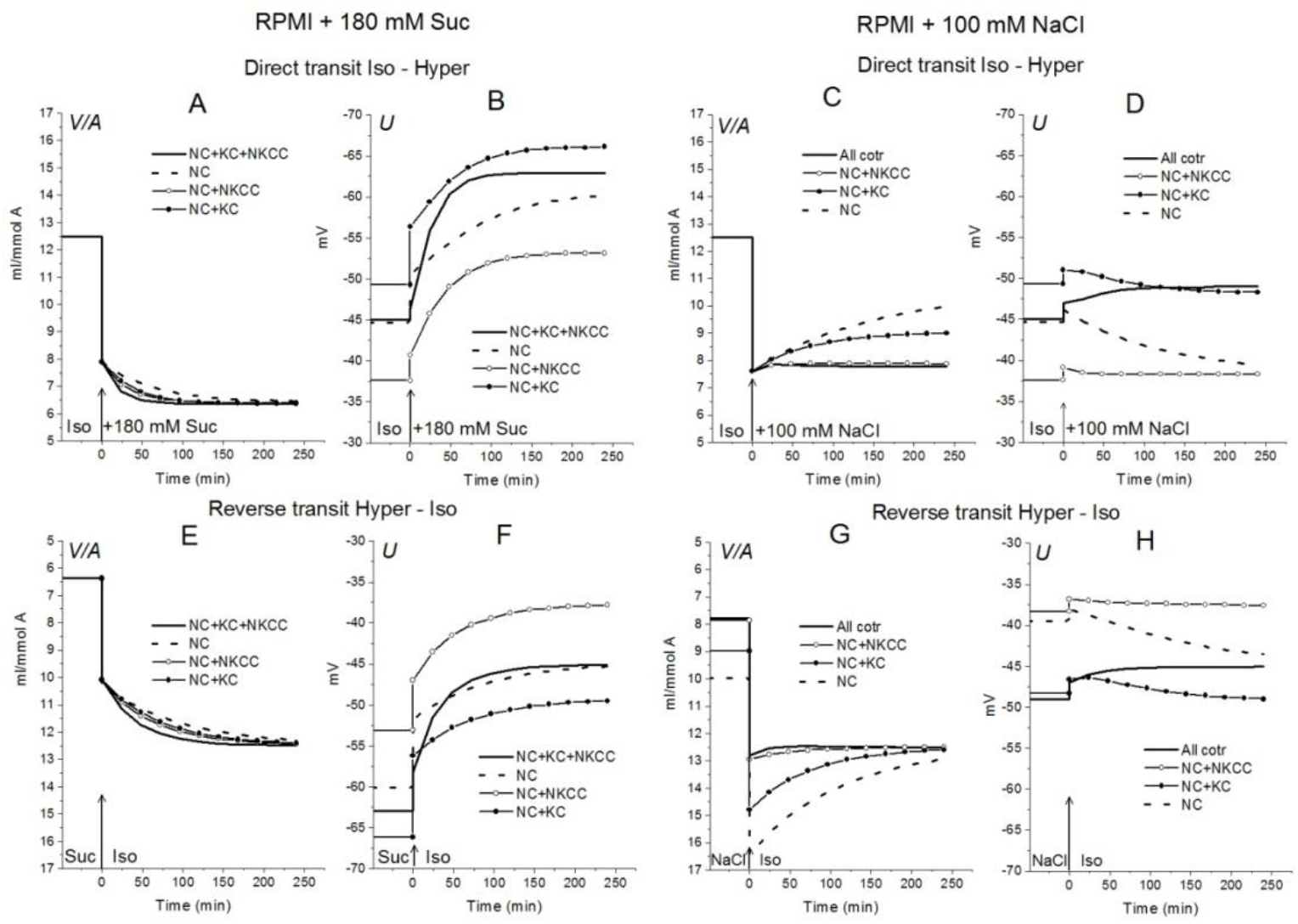
Rearrangement of ionic homeostasis caused by an increase in external osmolarity in the U937 cell model with different cotransporters and parameters like in cells balanced with the standard medium. Direct Iso-Hyper (**A-D**) and reverse Hyper-Iso (**E-H**) transitions.

### 2. Rearrangement of ionic homeostasis in hyperosmolar media at the simultaneous alteration of channels and transportersin cell model like U937

It is believed that specific changes in channels and transporters of the cell membrane are responsible for the regulation of cell volume under anisosmolar conditions, which are triggered by osmotic stress and can differ depending on the cell type (Hoffmann et al., 2009; Koivusalo et al., 2009; Hoffmann and Pedersen, 2010; Lang and Hoffmann, 2012; Jentsch, 2016; Pasantes-Morales, 2016; Delpire and Gagnon, 2018; Larsen and Hoffmann, 2020). This concept is based mostly on the studies of the effects of inhibitors and genetic cell modifications. Calculation of ionic homeostasis can provide a rational and more rigorous solution to the question of what changes in channels and transporters could underly the observed changes in ionic homeostasis under anisosmolar conditions. The parameters considered below as significant for RVI or AVD were determined in the simulation itself and taking into account opinions in the literature.

#### 2.1. Effect of NC cotransporter and Cl^-^ channels

Lew and Bookchin studying human reticulocytes were the first who showed that Na^+^ and Cl^-^ coupled transport across the cell membrane (NC) is an indispensable in quantitative description of the monovalent ion flux balance in cell (Lew et al., 1991). Analysis of the balance of ion fluxes in the main types of animal cells: the cells with low and high membrane potential and with high and low potassium content, led us to the conclusion that the K^+^/Na^+^ ratio and membrane potential depend primarily on the ratio of the Na^+^ and K^+^ channels permeability to the rate coefficient of the sodium pump, while the water content in the cell and intracellular Cl^-^ is mainly determined by the cotransporters and the permeability of the Cl^-^ channels (Vereninov et al., 2014; Yurinskaya et al., 2019; see also Jentsch, 2016; Dmitriev et al., 2019). NC cotransport in ionic homeostasis in U937 cells and, as we believe, in proliferating cells of a similar type, is the most important driver for active transport of Cl^−^ into cells and generation of a difference in electrochemical potential of Cl^−^ on the plasma membrane. A 3-fold increase in *inc* in the U937 cell model under standard isosmolar conditions increases the water content in cells from 12.5 to 19.64 ml/mmol *A*, and a 0.2-fold decrease decreases to 10.2 ml/mmol *A* (**Figure 3A).**

**Figure 3.**
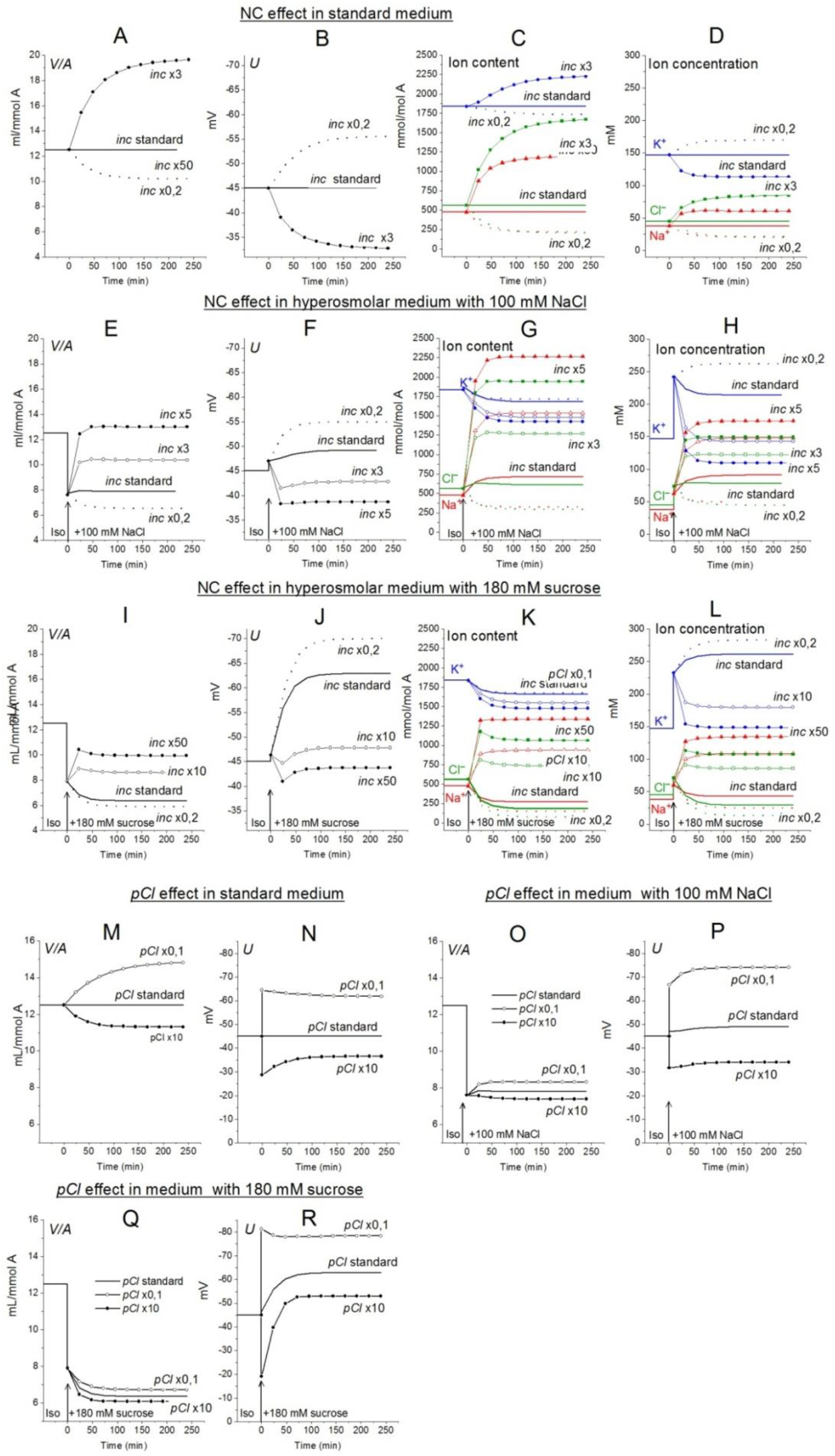
The effects of NC rate coefficient (**A-L**) and permeability coefficient of Cl^-^ channels (**M-R**) on the ionic homeostasis in U937 cell model under standard conditions and after transition to hyperosmolar medium with 180 mM sucrose or 100 mM NaCl. The calculation was carried out for a model with a full set of cotransporters. Changes in NC rate coefficient (*inc*) and in permeability coefficient of Cl^-^ channels (*pCl*) are shown in the graphs, other parameters remained unchanged and are shown in Table 2.

Small RVI occurs in a hyperosmolar NaCl medium even without any changes in *inc* or other membrane parameters. An increase in *inc* increases the RVI, while a decrease, on the contrary, causes a decrease in volume, like AVD. An increase in *inc* by about 5 times returns cell volume in the NaCl hyperosmolar medium to its level in the standard medium (**Figure 3E**). The effect of increasing *inc* in the sucrose hyperosmolar medium differs significantly from that in the NaCl hyperosmolar medium. A decrease in cell volume instead of an increase occurs in the sucrose hyperosmolar medium without changing *inc* (**Figure 3I, Table 6**). RVI can be obtained in the sucrose hyperosmolar only at a very significant increasing *inc* of about 10-50 times (**Figure 3I, Table 6**). The difference in the effects of *inc* in the hyperosmolar media with addition of sucrose and NaCl is due to the difference in changes of the electrochemical potential difference driving ions via NC pathway in the compared cases **(Figure 1F, L).**

The Cl^-^ channels and KC cotransporter in cells like U937 are the main antagonists of NC cotransporterbecause this is a significant pathway for the net Cl^-^ flux down electrochemical gradient (**Table 4,** yellow lines “Before”). The pair NC cotransporter-Cl^-^ channels is a powerful regulator of the cell water balance. This is since active transport of Cl^-^ into the cell via NC, which increases its intracellular concentration above the equilibrium level, is equivalent to an increase in the amount of internally sequestered Donnan anions which causes an increase in the water content in the cell. In the considered U937 cell model the effect of pCl variation on ion homeostasis is less significant than the effect of NC. For example, the 10-fold decrease of pCl causes an increase in cell volume by a factor 1.18 (from 12.5 to 14.83 ml/mmol *A*, **Figure 3M**) in the standard medium and, respectively, 1.09 (from 7.6 to 8.32 ml/mmol *A*) in hyperosmolar medium with the addition of 100 mM NaCl (**Figure 3Q)**. The 3-fold increase in NC rate coefficient turns out to be stronger than 10-fold decrease in the permeability of Cl^-^ channels. This is because there are other pathways for downhill movement of Cl^-^ besides channels in the cells like U937. The channel part of the Cl^-^ downhill net flux is only about half; the other half is the flux through the KC pathway and about 0.6 % is the flux through the NKCC cotransporter (**Table 4,** yellow lines “Before”).

#### 2.2. Effect of NKCC and KC cotransporters (ikc, inkcc)

The specifically weak effect of NKCC on ion homeostasis of cells balanced with the normal medium was discussed in the previous section.This is due to a small integral electrochemical difference driving ions via NKCC under normal conditions. Accordingly, the net fluxes of K^+^, Na^+^ and Cl^-^ via NKCC in cells like U937 balanced with normal isosmolar medium are small and increase when cells are transferred into the hyperosmolar medium (**Table 4,** compare the lines “Before” marked in yellow with lines marked in green). Changes in *inkcc*, simultaneously with an increase in external osmolarity, change both the dynamics of the transition and homeostasis in a new state of balance in a hyperosmolar medium with the addition of NaCl, but not in a sucrose medium (**Figure 4)**. However, even in the NaCl hyperosmolar medium, a 10-fold increasing or decreasing *inkcc* affects the new balanced cell volume by no more than 1.16 times. A decrease in *inkcc* by a factor 0.1 causes RVI in the NaCl hyperosmolar medium, while its increase causes volume decrease like AVD. A decrease in the KC rate coefficient, *ikc*, affects the water balance in NaCl hyperosmotic medium in a similar way as NKCC, but its effect on membrane potential is stronger than that of NKCC (**Figure 4I, J**).

**Figure 4.**
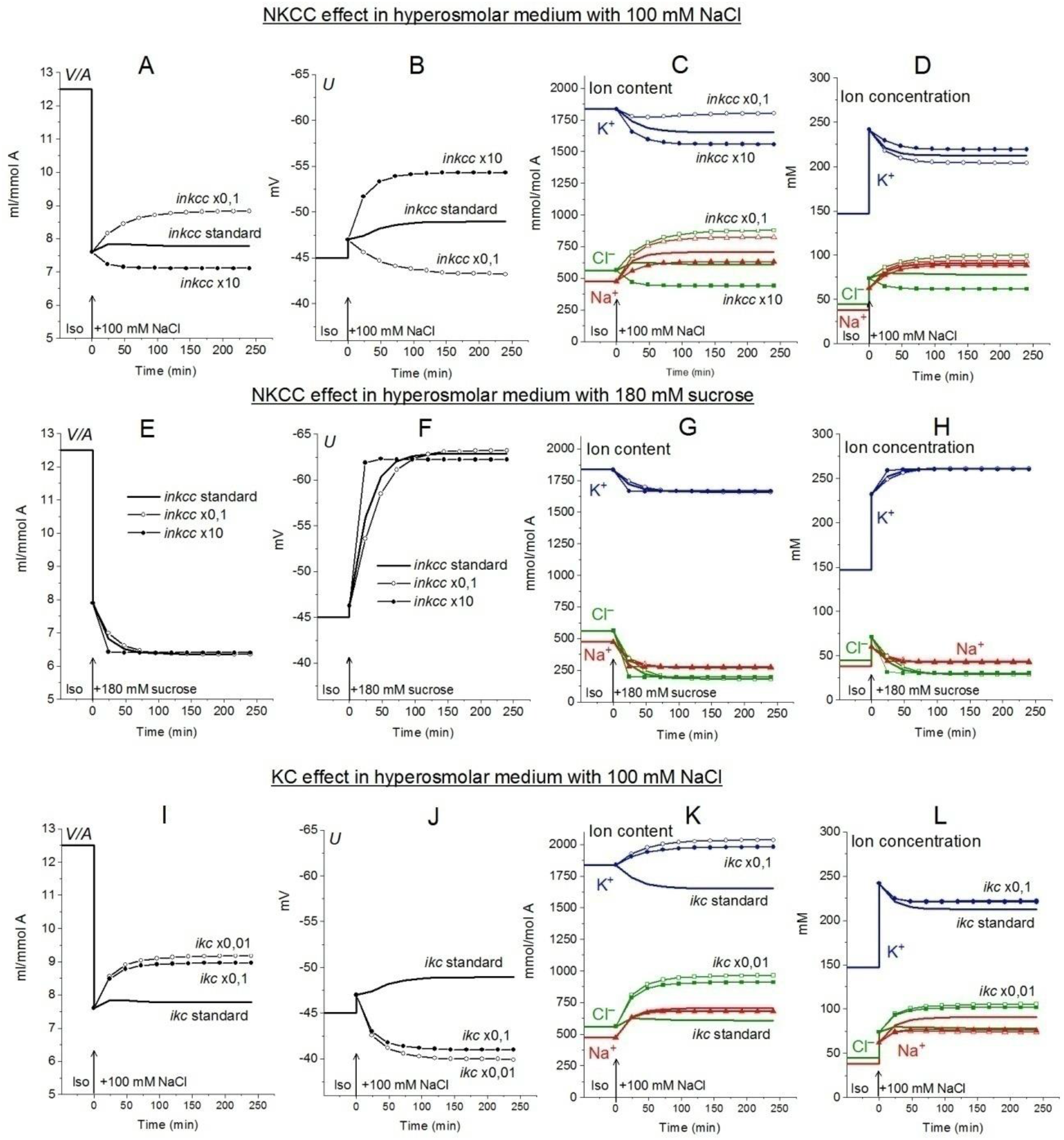
Dependence of ionic homeostasis in U937 cell model during transition to hypertonic medium with 100 mM NaCl (**A-D**, **I-L**) or 180 mM sucrose (**E-H**) on the rate coefficients *inkcc* (**A-H**) and *ikc* (**I-L**), changing simultaneously with external osmolarity. The calculation was carried out for a model with a full set of cotransporters. Changes in NKCC and KC rate coefficients are shown in the graphs, other parameters remain unchanged and are shown in Table 2.

#### 2.3. Effect of Na^+^ channels and non-selective hypertonicity-induced cation channels, HICC

In view of the experimental data indicating that Na^+^ channels and the channels known as HICC (Wehner et al., 2003, 2006; Plettenberg et al., 2008) in some cell types can play a role in RVI (Hoffmann et al., 2009) the possible effects of these channels were examined in U937 cell model. The Na^+^, K^+^ non-selective channels HICC were mimicked by increasing pNa 10 times with simultaneous increasing pK 1.6 times. This is equivalent to the addition of the HICC in an amount 1.6 times higher than the standard number of K^+^ channels. It turned out that an increase in pNa upon transition to a hyperosmolar medium causes RVI in the considered system, but only in a hyperosmolar medium with the addition of NaCl (**Figure 5**). No RVI occurs in case of the medium with 180 mM sucrose. An increase in pNa in cells placed in a standard 310 mOsm medium causes a much greater increase in cell volume than in a hyperosmolar medium with additional NaCl under the same conditions. The effect of increasing pNa alone is attenuated in case of HICC which include increasing pK. This is due to the opposite effect of changes in pNa and pK on ionic homeostasis.

**Figure 5.**
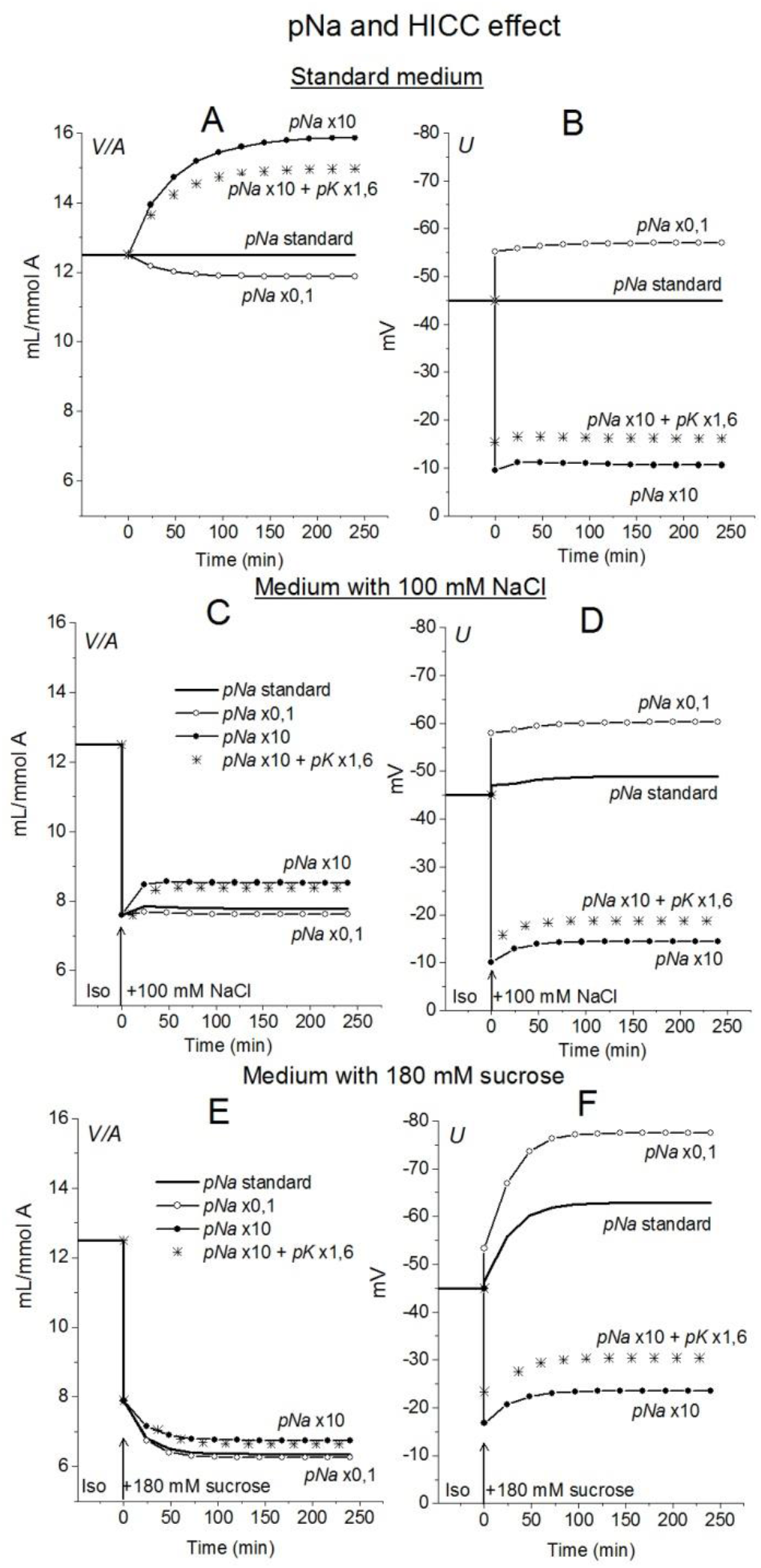
The effect of changes in the Na^+^ and HICC channel permeability coefficients on ion homeostasis in the U937 cell model under standard conditions (**A, B**) and in a hyperosmolar medium with 100 mM NaCl (**C, D**) or 180 mM sucrose (**E, F**). The calculation was carried out for a model with a full set of cotransporters. Changes in *pNa*, and HICC (*pNa+pK*) are shown in the graphs, other parameters remained unchanged and are shown in Table 2.

The examples in **Figures 3–5** demonstrate which changes in membrane channels and transporters of the cell in a hyperosmolar medium can be associated with RVI, and which, on the contrary, with AVD. **Table 5** summarizes the main differences associated with changes in the new balanced state in hyperosmolar media under various conditions. One of the intuitively unpredictable points is that the same changes in channels and transporters, for example, a decrease in *ikc* or *pCl*, can cause RVI (marked in yellow in **Table 5**) in a hyperosmolar medium with added NaCl and oppositely directed AVD (marked in blue) in a hyperosmolar medium with sucrose. It turned out that in cells such as U937, there is only one way to obtain RVI in the hyperosmolar medium with sucrose by changing the membrane parameters. This is an increase in parameter *inc*. There are more variants in the hyperosmolar medium with added NaCl. One of the intuitively unpredicted points is that the balanced state in hyperosmolar medium with added NaCl is associated mostly with an increase in cell Na^+^ content whereas K^+^ content decreases. The only exception is when the RVI is caused by a strong decrease in the rate coefficient *ikc*. In the hyperosmolar sucrose medium, the balanced state of cells is characterized, for the most part, by a stronger decrease in the Na^+^ content than in K^+^. Only an increase in *inc, HICC* and a decrease in *Pump beta* cause a stronger decrease in K^+^ and an increase in the Na^+^ content in the balanced state.

**Table 5.**
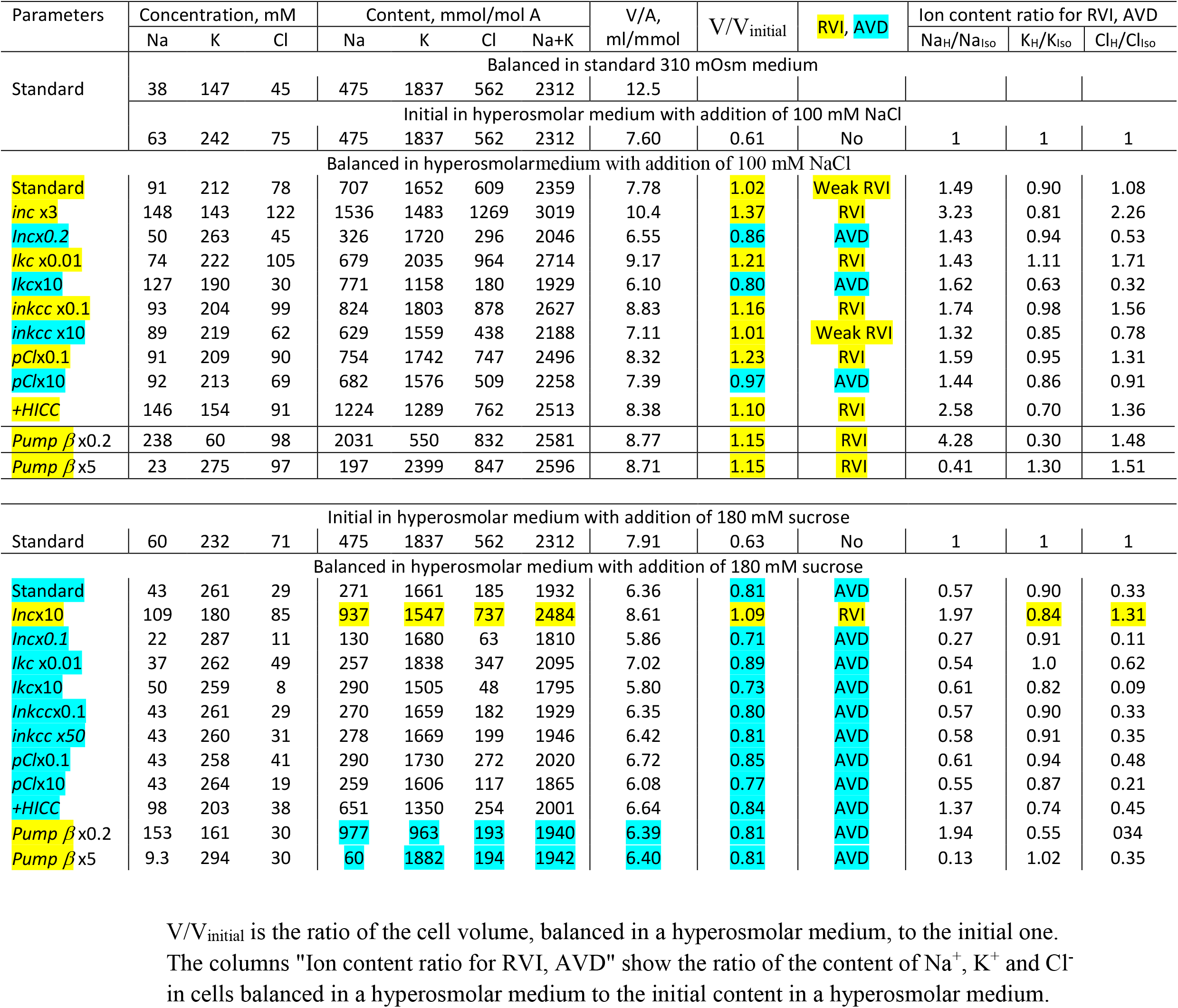
Dependence of RVI and AVD in the U937 cell model on the rate coefficients *inc*, *ikc*, *inkcc, pCl*, the permeability coefficient of the *HICC* channel (*pNa+pK*), and pump rate coefficient *β* changing simultaneously with an increase in external osmolarity. Other parameters remain unchanged. The calculation was carried out for a model with all cotransporters. hp = 240 in all cases except hp=800 for *Pump β*. The RVI effect is marked in yellow, the AVD effect is marked in blue.

### 3. Changes in water and ionic balance in living cells such as U937 in a hyperosmolar media. Dual responseoflivingcellsto hyperosmolar challenge

The response of living cells to transfer in hyperosmolar medium is more complicated than in the electrochemical model since the properties of membrane channels and transporters can be changed by physically unpredictable way through the intracellular signaling network. Cell shrinkage in hyperosmolar medium triggers two complex general cellular responses, which are characterized by the opposite direction of volume change and develop with a shift in time (Yurinskaya et al., 2011, 2012, 2017). This is the AVD associated with apoptosis (Okada et al., 2001), and the oppositely directed RVI, which precedes the AVD (Yurinskaya et al., 2012). In our experience, the analysis of the distribution of cells in the density gradient is the best method for separating the primary rapid physical decrease in the volume of cells in the hyperosmolar environment and the specific processes of RVI and AVD (**Figure 6**). Due to the time shift, RVI and AVD can be observed on the same sample of cells. An essential detail is that the transition from RVI to AVD in the cell population manifests itself as a change in the ratio between the number of cells in the RVI and AVD stages. The number of RVI cells decreases over time and the number of AVD cells increases (**Figure 6M**). This indicates that the transition from RVI to AVD in each cell is fast. Ionic changes underlying RVI and AVD in hyperosmolar media, obtained on K562, Jurkat, and U937 cells in another separate series of experiments, where all these cell types were studied simultaneously, are presented in **Figure 7** and **Table 6**.

**Figure 6.**
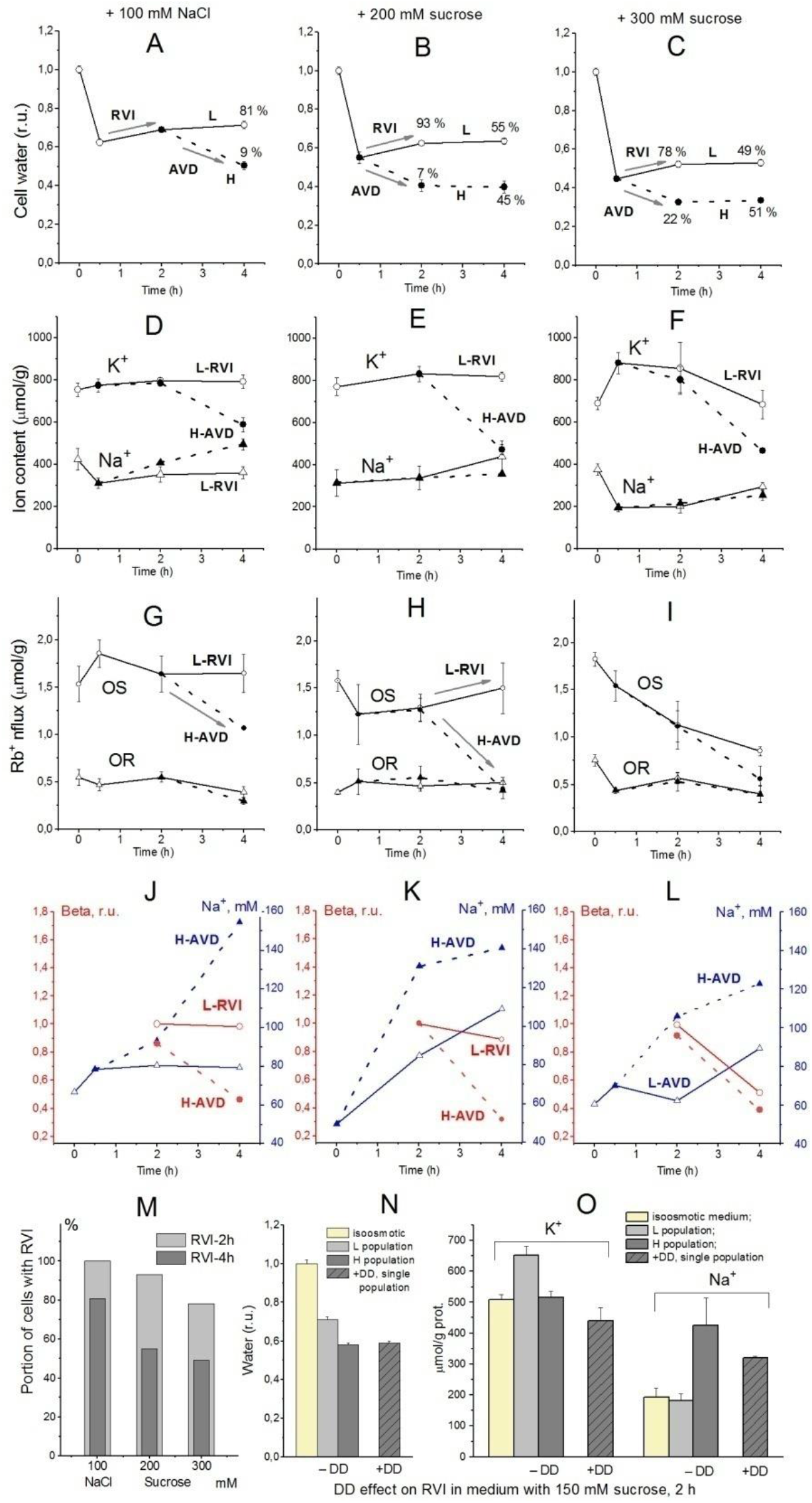
Effect of hyperosmolar medium on living U937 cell. (**A-C**) cell water content, (**D-F**) intracellular K^+^, Na^+^ content, (**G-I**) ouabain-sensitive (OR) and –resistant (OR) Rb^+^ influxes, (**J-L**) Na^+^ concentrations and *beta*, (**M**) the percentage of cells with RVI (fraction L) after 2-hour and 4-hour incubation in a hypertonic medium assessed by protein, (**N, O**) DMA+DIDS (DD) effect on RVI in hyperosmolar medium with 150 mM sucrose. Solid lines with open symbols indicate light (L) cell subpopulation going RVI stage; dotted lines with filled symbols indicate heavy (H) cell subpopulation going AVD stage. Data at time zero represent cells in normal RPMI medium. Mean ± SEM values were calculated from at least 3 independent experiments. (**N, O**) DMA (0.05 mM) and DIDS (0.5 mM) were added simultaneously with addition of 150 mM sucrose. FromYurinskaya et al., 2012 modified.

**Figure 7.**
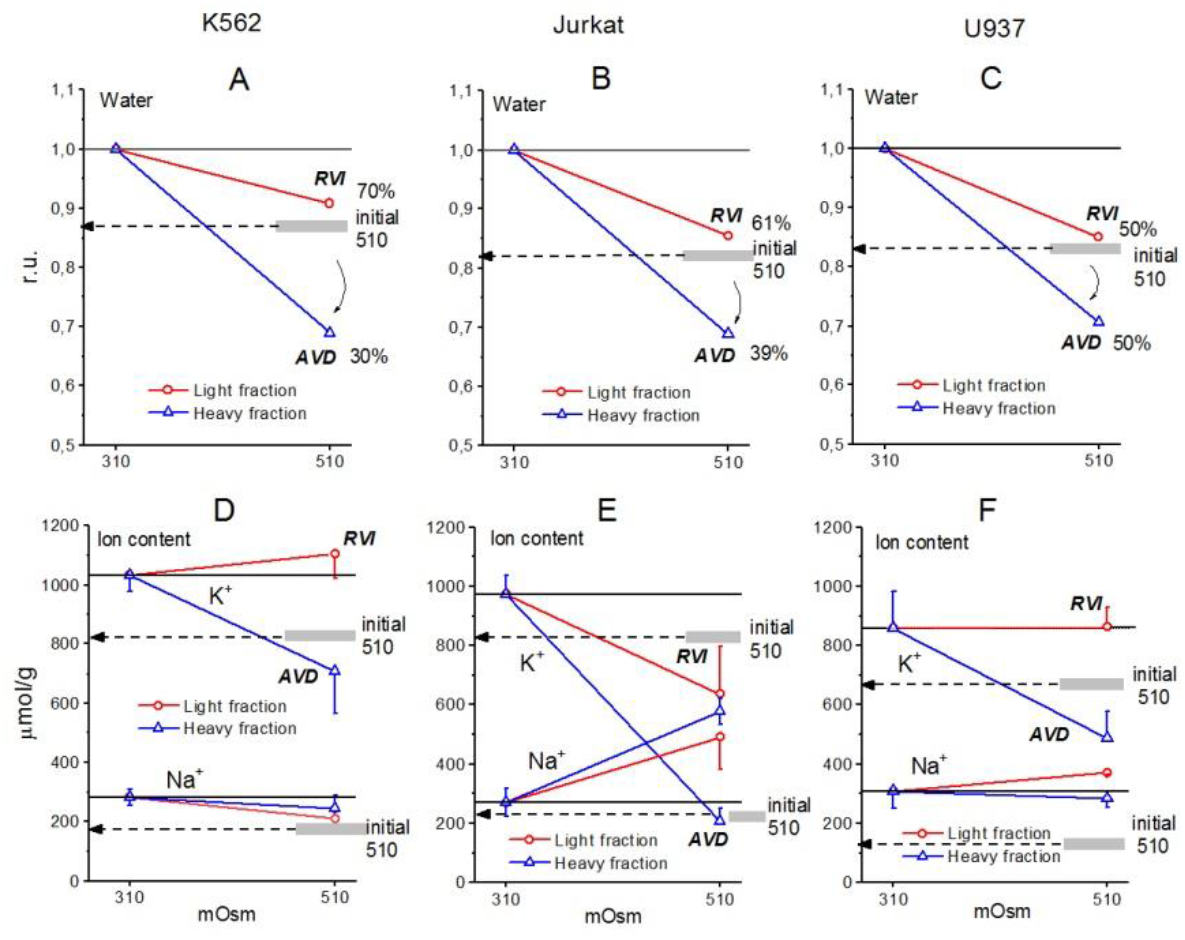
Cell water, K^+^, and Na^+^ content in living K562 (**A, D**), Jurkat (**B, E**), and U937 (**C, F**) cells before and after 4 h incubation in hyperosmolar medium with 200 mM sucrose (510 mOsm). The broad gray lines show the level of the initial water and ion content in hyperosmolar medium (15 min incubation).

**Table 6.**
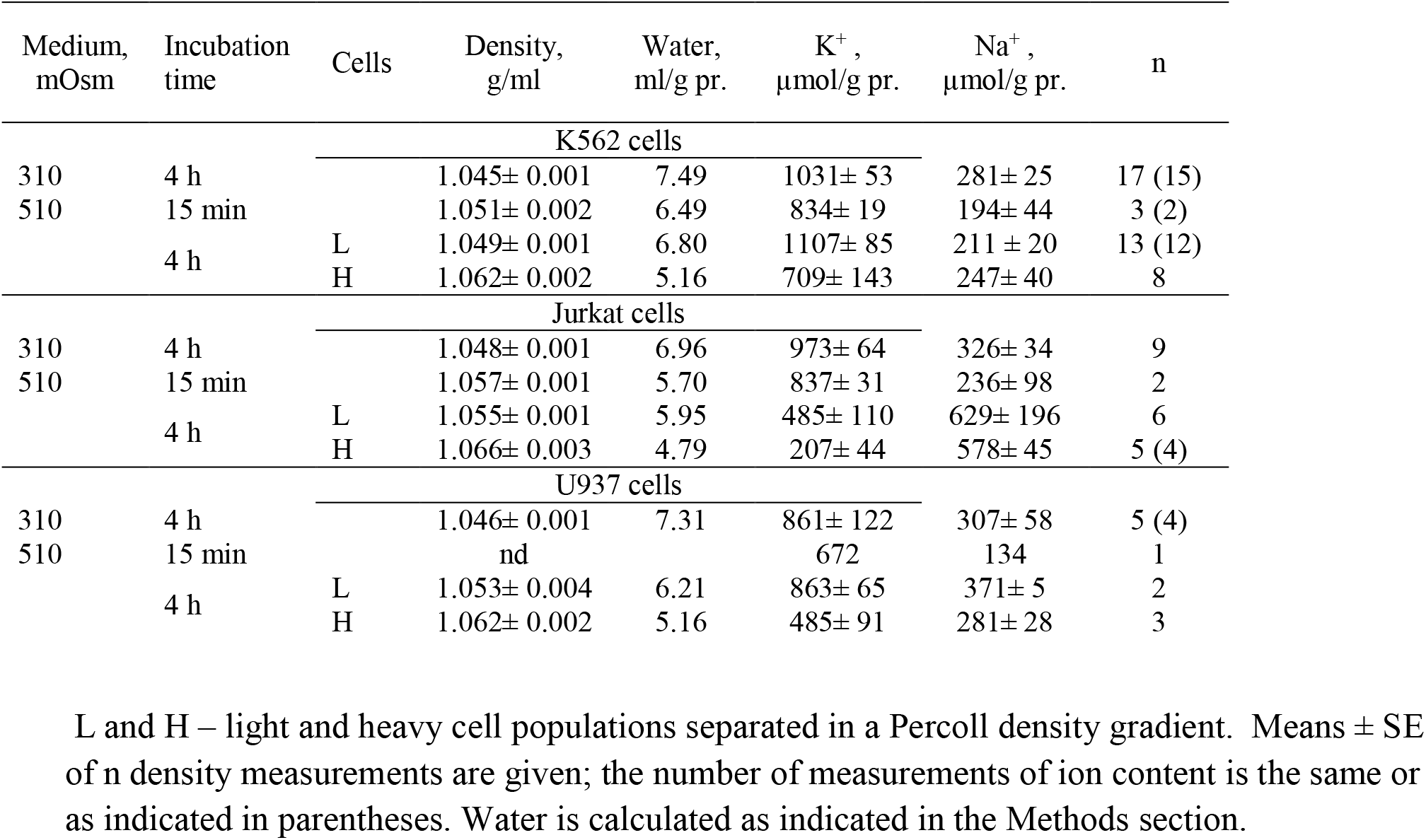
Changes in the buoyant density, K^+^ and Na^+^ content in living K562, Jurkat, and U937 cells transferred to a hyperosmolar medium with adding 200 mM sucrose.

The RVI and AVD mechanisms are discussed in more detail below, considering computer simulations of the system and data related to U937 cells (Yurinskaya et al., 2011, 2012, 2017). In the experiments shown in **Figure 6**, the mean values of ions and water during the first 2 hours characterize mainly cells at the RVI stage. At the time point 4 h the light, RVI, and heavy, AVD, subpopulations could be analyzed separately. According to modeling in a hyperosmolar medium with sucrose, RVI in U937 cells is possible only with an increase in the coefficient *inc* (**Table 5).** This should be associated with an increase in the intracellular content of Na^+^ and Cl^-^ and a slight change in the content of K^+^. It is these changes in the content of Na^+^ and K^+^ that were observed in experiments with U937 cells for a light subpopulation of RVI in a hyperosmolar medium with sucrose (**Figure 6E, Table 6**). The key role of the NC cotransporter in RVI is independently confirmed by the blocking effect of the combination of dimethylamiloride, a known inhibitor of Na/H exchanger, and DIDS, which inhibits the Cl/HCO_3_ exchanger (**Figure 6N, O**). In hyperosmolar medium with addition NaCl an increase in *inc* or a decrease in *ikc, inkcc, pCl* or appearance of channels *HICC* increases RVI (**Figures 3E, O, 4A, I, Table 5).** In all these cases, RVI is associated with an increase in the total content of K^+^ and Na^+^, but the relative changes in the content of K^+^ and Na^+^ depend on which parameter changes. In living U937 cells, RVI in a hyperosmolar medium with the addition of NaCl is small, the content of K^+^ and Na^+^ changes insignificantly (**Figure 6A, D**). However, the available data are insufficient to differentiate the mechanism of RVI in these cases in more detail.

In experiments where cells U937, K562, and Jurkat were studied in parallel in hyperosmolar medium with 200 mM sucrose for 4 h the certain differences between cell species were revealed (**Table 6, Figure 7**). RVI at the time point 4 h was rather small but differences in K^+^ and Na^+^ content in K562, Jurkat and U937 cells were significant. Further research is needed to better understand these cell-species differences.

AVD in living U937 cells incubated in a hyperosmolar sucrose medium is slightly stronger than in a medium with additional NaCl of almost the same osmolarity. In both cases, AVD is associated with a significant decrease in the K^+^ content in cells, while the Na^+^ content in cells increases in a hyperosmolar NaCl medium and changes insignificantly in a sucrose medium. This agrees with the model prediction. It was shown earlier that a decline in ouabain-inhibitable Rb^+^ (K^+^) influx due to a decrease in the pump rate coefficient, *beta*, plays significant role in AVD during apoptosis of U937 cells induced with staurosporine (Yurinskaya et al., 2019, 2020). AVD in U937 in hyperosmolar media in experiment described in the present article is also associated with a decrease in the OS Rb^+^ influx whereas the decline in OR influx is much less (**Figure 6G-L,** H-AVD subpopulation). Comparison of the changes in OS Rb^+^ influx with the changes in concentration of Na^+^ in cell water indicates that a decline in the pump Rb^+^ influx manifests mostly the changes in the pump rate coefficient, i.e., in intrinsic pump properties. There are no significant changes in the in OS Rb^+^ influx and the pump rate coefficient in cells of the light subpopulations in 100 mM NaCl and 200 mM sucrose hyperosmolar media (**Figure 6G, H, J, K**; L-RVI subpopulation).

## Discussion

This work continues our studies aimed at developing a modern mathematical description of a complex interrelationship of monovalent ion fluxes via all main parallel pathways across the cell membrane, including all cation-chloride cotransporters, which all together determine the entire water and ion balance in animal cell. Earlier this mathematical description was applied to cases of blockage of the Na/K pump (Vereninov et al., 2014, 2016; Yurinskaya et al., 2019, Yurinskaya and Vereninov, 2021), apoptosis (Yurinskaya et al., 2019, 2020), and the response to hypoosmolar stress (Yurinskaya and Vereninov, 2021). Now it is used to analyze the cellular response to hyperosmolar stress, during which two opposite effects of this stress on water balance, RVI and AVD, occur in the same cells, following each other with a delay.

As in the pioneering fundamental works (Jakobsson, 1981; Lew, Bookchin 1986; Lew et al., 1991; Lew, 2000), our description is based on the classification of ion transport pathways according to the acting driving forces and the type of coupling of the fluxes of various ions and on characterization of the kinetics of ion transfer using a single rate coefficient. This makes the description independent of the specific molecular mechanism of ion movement via channels and transporters. Only such a holistic approach enables quantitative analysis and allows to predict real changes in cell ion and water balance and electrochemical ion gradients on the cell membrane. Due to complex interdependence of ion fluxes related to parallel pathways the individual unidirectional fluxes along the separate channels or transporter usually cannot be measured directly. They must be calculated considering all fluxes in the cell. Since we cannot measure individual fluxes, we had to use alternative approach to validateof the mathematical description. The validation was based on the analysis of the cell system dynamics by stopping sodium pump. Calculation of the unidirectional fluxes is important for studying the functional expression of separate channels and transporters using specific inhibitors because enables to determine when the use of inhibitors can reveal fluxes related to specific channels or transporters, and when not, due to masking by fluxes across parallel pathways.

Our description is applied to experimental data obtained from U937 cells cultured in suspension, which allows a wide range of assays to be used without cell change caused by isolation, and includes cell water determination using buoyant density, cell ions using flame photometry, and optical methods using flow cytometry. In recent years, in neurobiology and other fields, there has been a growing interest in disorders of ionic and water homeostasis of cells (Dijkstra et al.., 2016; Pasantes-Morales, 2016; Casula et al., 2017; Wilson and Mongin, 2018; Bortner and Cidlowski, 2020; Van Putten et al., 2021). However, in many cases of practical importance cells cannot be isolated for proper assays. Therefore, U937 cells can serve a useful model for understanding the general mechanisms of cell water and ionic balance regulation.

An essential part of the results is a developed software supplied with executable file that allows one to determine the role of each type of cotransporters or channel in the regulation of the ionic and water balance of cells in the context of the cell type and actual conditions. The variety of the effects caused by changing channels and transporters is vast, even for one type of cells, as shown by the example of U937 cells demonstrated in previous and present studies.Even the limited number of examples selected to illustrate our approach took up a lot of space.It is clear that serious research in this area is impossible without calculating a specific system.

Computation of the possible changes in ionic and water balance in the U937 cell model specifically in hyperosmolar media has revealed many interesting and, at first glance, unexpected things. (1) An AVD-like effect can occur in a sucrose-supplemented hyperosmolar medium and an RVI-like effect in a NaCl-supplemented hyperosmolar medium without altering membrane channels and transporters due to time-dependent changes in the forces moving monovalent ions across the cell membrane. It is noteworthy that this is observed only with some types of cotransporters. (2) Changes in the cell membrane potential in hyperosmolar media of both types significantly depend on the set of cotransporters, despite their “electroneutrality”, as predicted by more general considerations (Stanton, 1983). (3) The sign of the forces driving ions through the NC, KC, and NKCC cotransporters under certain conditions is of paramount importance for the role of these cotransporters in the regulation of the ionic and water balance of the cell during RVI and AVD. Simulation draws attention to the analysis of changes in driving forces in addition to changes in channel and transporters properties.

Study of living U937 cells shows that RVI and AVD responses to the hyperosmolar medium are caused not only by changes in ion channels and transporters but, in addition, by the redistribution of organic osmolytes, regulated by signals from the intracellular signaling network. A similar redistribution of organic osmolytes has been shown for many other cells (see reviews: Kirk, 1997; Lambert et al., 2008; Hoffmann et al., 2009; Koivusalo et al., 2009; Pasantes-Morales, 2016). Modeling can tell nothing about which intracellular signals change channels and transporters or change the intracellular content of impermeant osmolytes or their charge.However, it is possible to recognize and estimate quantitatively the alteration of executing mechanisms regulating cell ion and water balance.

In the case of living cells, such as U937, when the required minimum of experimental data for calculations is available, our results show that RVI in a hyperosmolar medium with sucrose is possible only due to an increase in the coefficient *inc*. In this case, RVI should be associated with an increase in the intracellular content of Na^+^ and Cl^-^ and a slight change in the content of K^+^. It is these changes in the content of Na^+^ and K^+^ that were observed in experiments with U937 cells for a light subpopulation of RVI in a hyperosmolar medium with 200 mM sucrose. The key role of the NC cotransporter in RVI is independently confirmed by the blocking effect of the combination of dimethylamiloride, a known inhibitor of Na/H exchange, and DIDS, which inhibits the Cl/HCO_3_ exchange.

In a hyperosmolar medium with the addition of NaCl, a slight increase in cell volume with time, like RVI, can occur in accordance with modeling even without changing the membrane parameters in a cell with NC or NC+KC cotransporters, but without NKCC. In this medium, an increase in *inc* or a decrease in *ikc, inkcc, pCl* or appearance of channels *HICC* increases RVI.In all these cases, RVI is associated with an increase in the total content of K^+^ and Na^+^, but the relative changes in the content of K^+^ and Na^+^ depend on which parameter changes.Living U937 cells in a hyperosmolar medium supplemented with NaCl show a small RVI and insignificant changes in the content of K^+^ and Na^+^. However, the available data are insufficient to differentiate in more detail the mechanism of RVI in these cases.

The AVD response demonstrated by a heavy subpopulation of living U937 cells incubated in a hyperosmolar medium with sucrose is slightly stronger than in a medium supplemented with NaCl of almost the same osmolarity.In both cases, AVD is associated with a significant decrease in the K^+^ content in cells, while the Na^+^ content in cells increases in a hyperosmotic NaCl medium and changes insignificantly in a sucrose-containing medium. This is consistent with the prediction of the model.It was shown earlier that a decrease in the ouabain-inhibited Rb^+^ (K^+^) influx due to a decrease in the pumping rate coefficient beta plays an important role in AVD during apoptosis of U937 cells induced by staurosporine (Yurinskaya et al., 2019, 2020). AVD in U937 in hyperosmolar media in the experiments described in the present article is also associated with a decline in the pump Rb^+^ influx due to mostly the changes in the pump rate coefficient, i.e., in intrinsic pump properties. No significant changes in the OS Rb^+^ influx and the pump rate coefficient were observed in cells of the light subpopulations in 100 mM NaCl and 200 mM sucrose hyperosmolar media demonstrating the RVI response.

The main conclusion of this study, which demonstrates an example of analysis of the mechanism of the RVI and AVD responses to hyperosmolar stress of cells such as U937, is that computer calculations are an indispensable tool for studying mechanisms not only of RVI and AVD, but all phenomena associated with the regulation of the entire electrochemical system of the cell.

## Supporting information

How to Use BEZ02BC

Input data

The executable file

## DATA AVAILABILITY STATEMENT

The original contributions presented in the study are included in the article/Supplementary Material, further inquiries can be directed to the corresponding author.

## AUTHOR CONTRIBUTIONS

Both authors contributed to the design of the experiments, performed the experiments, analyzed the data, and approved the final version of the manuscript and agreed to be accountable for all aspects of the work. Both persons designated as authors qualify for authorship.

## ACKNOWLEDGMENTS

We are grateful to Dr. Igor A. Vereninov for correcting the manuscript and suggestions for improvement. Our thanks to Igor Raikov, a student of the Alferov Federal State Academic University RAS, Russia, for checking the use of the BEZ02BC file on a 32-bit computer.

## SUPPLEMENTARY MATERIAL

Executable file to the programme code BEZ02BC and Instruction: How to use programme code BEZ02BC.doc are attached to the article electronic version.

