## Supplementary material for "Role of cation-chloride cotransporters, Na/K-pump, and channels in cell water/ionic balance regulation under hyperosmolar conditions: in silico and experimental studies of opposite RVI and AVD responses of U937 cells to hyperosmolar media": How to Use BEZ02BC

How to use the executable file for the program BEZ02BC.

1. The executable file BEZ02BC (BEZ02BC.doc when e-mailed) is best used on a 32-bit

computer with Windows OS. The user should in this case:

a. Locate files DATAB.doc and BEZ02BC.doc in the same folder.

b. Change the extension of BEZ02BC from “doc” to “exe” (do not try to open BEZ02BC.doc. It is unreadable!) and the extension DATAB.doc to .txt.

c. Check the DATAB, the file must correspond to the selected parameters and concentrations, click "Save" (simple “DATAB.txt” without specific name)

d. Run the executable file BEZ02BC….. and wait until the process is completed and the file RESB.txt appears.

e. Rename and save the obtained file RESB.txt because in a new running cycle it will be lost.

RESB files can be easily imported by ORIGIN or another program for further processing (the asterisks must be retained).

f. The displayed values of fluxes as well as OSOR correspond to the latest time point. The values of fluxes for other moments can be obtained by setting the necessary time interval with the *hp* value. It is necessary to perform several calculation cycles with a series of the corresponding *hp* to obtain the time course of the fluxes.

g. Some readers of our previous publications have expressed doubt that using our tool it is possible to obtain a unique set of parameters that provide an agreement between experimental and calculated data. Our mathematical comments on this matter can be found in Yurinskaya et al., 2019, P.12.

2. To run the executable file on a 64-bit machine, the following additional steps should be taken:

h. Download the School Pak package via Internet and run Norton Commander (NCD).

i. Set in NCD the same folder as the folder in Windows where the DATAB.txt and executable file BEZ02BC are located.

j. Correct the file DATAB.txt if necessary, in the Windows folder.

k. Run executable file in the NCD folder and read RESB.txt in the analogous Windows folder.

Several DATAB and appropriate RESB options are presented below as examples:

Example 1 “Iso standard 240”, Cells with all cotransporters:

DATAB

na0	k0	cl0	B0	kv	na	k	cl	beta gamma

140.0	5.8 116.0 48.2 1.0 38.0 147.0 45.0 0.039 1.50

pna	pk	pcl	inc	ikc	inkcc	hp	kb

0.00170 0.01150 0.01100 0.000070 0.0000800 0.0000000080 240 0.0

RESB

t	U	na	k	cl	V/A	mun	muk	mucl	naC	kC	clC

0 -45.0 38.0 147.0 45.0 12.50 -79.9	41.3	19.8	475.0	1837.5	562.5

24 -45.0 38.0 147.0 45.0 12.50 -79.9	41.3	19.8	475.0	1837.5	562.5

....................................................................................

240 -45.0 38.0 147.0 45.0 12.50 -79.9	41.3	19.8	475.0	1837.4	562.4

*	na0	k0 cl0	B0	kv	na	k	cl	beta gamma

*	140.0	5.8 116.0 48.2 1.000	38.0 147.0	45.0 0.039 1.50

*	pna	pk	pcl	inc	ikc	inkcc	hp	kb

*	0.00170 0.01150 0.01100 0.0000700 0.0000800 0.0000000080	240 0.000000

*	Net_flux PUMP	Channel	NC	KC	NKCC

*	Na -1.4820 0.4679 1.0171 0.0000 -0.0031

*	K	0.9880 -0.5096 0.0000 -0.4753 -0.0031

*	Cl	0.0000 -0.5357 1.0171 -0.4753 -0.0061

*	Influx	PUMP	IChannel INC	IKC	INKCC

*	Na	0.0000 0.4927 1.1368 0.0000 0.0874

*	K	0.9880 0.1381 0.0000 0.0538 0.0874

*	Cl	0.0000 0.4889 1.1368 0.0538 0.1748

*	Efflux	PUMP	EChannel ENC	EKC	ENKCC

*	Na -1.4820 -0.0248 -0.1197 0.0000 -0.0905

*	K	0.0000 -0.6477 0.0000 -0.5291 -0.0905

*	Cl	0.0000 -1.0246 -0.1197 -0.5291 -0.1809

*	z	OSOR (A/V)*1000

*	-1.75 3.54	80.00

Example 2 “Hyper 100 mM NaCl Cells with all cotr.”:

DATAB

na0	k0	cl0	B0	kv	na	k	cl	beta gamma

240.0	5.8 216.0 48.2 1.645	38.0 147.0 45.0 0.039 1.50

pna	pk	pcl	inc	ikc	inkcc	hp kb

0.00170 0.01150 0.01100 0.000070 0.0000800 0.0000000080 240 0.0

RESB

t	U	na	k	cl	V/A	mun	muk mucl	naC	kC	clC

0 -47.0 62.5 241.8 74.0 7.60 -82.9	52.6	18.4	474.8	1836.8	562.3

24 -47.4 81.3 221.5 79.7 7.84 -76.3	49.8	20.8	637.3	1736.8	624.7

48 -48.2 87.4 215.4 79.5 7.83 -75.1	48.4	21.5	684.5	1687.9	623.1

………………………………………………………………………………………………………………

168 -48.9 90.8 212.4 78.3 7.78 -74.9	47.2	21.9	706.3	1652.1	609.1

192 -49.0 90.8 212.4 78.3 7.78 -74.9	47.2	21.9	706.4	1651.9	608.9

216 -49.0 90.8 212.3 78.3 7.78 -74.9	47.2	21.9	706.4	1651.8	608.9

240 -49.0 90.8 212.3 78.3 7.78 -74.9	47.2	21.9	706.5	1651.7	608.9

*	na0	k0 cl0	B0	kv	na	k	cl	beta gamma

*	240.0	5.8 216.0 48.2 1.645	62.5 241.8	74.0 0.039 1.50

*	pna	pk	pcl	inc	ikc	inkcc	hp	kb

*	0.00170 0.01150 0.01100 0.0000700 0.000080 0.0000000080	240 0.0

Net_flux PUMP	Channel	NC	KC	NKCC

*	Na -3.5420 0.8366	3.1312 0.0000 -0.4257

*	K	2.3613 -0.7063	0.0000 -1.2295 -0.4257

*	Cl	0.0000 -1.0503	3.1312 -1.2295 -0.8514

*	Influx	PUMP	IChannel	INC	IKC	INKCC

*	Na	0.0000	0.8904	3.6288 0.0000 0.5196

*	K	2.3613	0.1456	0.0000 0.1002 0.5196

*	Cl	0.0000	0.8288	3.6288 0.1002 1.0391

*	Efflux	PUMP	EChannel	ENC	EKC	ENKCC

*	Na -3.5420 -0.0539 -0.4976 0.0000 -0.9453

*	K	0.0000	-0.8518	0.0000 -1.3297 -0.9453

*	Cl	0.0000	-1.8791 -0.4976 -1.3297 -1.8906

*	z	OSOR (A/V)*1000

*	-1.75 3.09 128.56

Example 3 “Hyper 180 mM Sucrose Cells with all cotr.”:

DATAB

na0	k0	cl0	B0	kv	na	k	cl	beta	gamma

140.0	5.8 116.0 228.2 1.58	38.0 147.0 45.0 0.039 1.50

pna	pk	pcl	inc	ikc	inkcc	hp kb

0.00170 0.01150 0.01100 0.000070 0.0000800 0.0000000080 240 0.0

RESB

t	U	na	k	cl	V/A	mun	muk mucl	naC	kC	clC

0 -46.3 60.0 232.3 71.1 7.90 -68.9	52.3	33.2	474.2	1834.6	561.6

24 -55.9 47.6 252.2 43.6 6.82 -84.6	44.9	29.7	324.8	1719.7	297.3

48 -60.3 44.0 258.4 34.0 6.51 -91.2	41.1	27.5	286.5	1681.8	221.1

………………………………………………………………………………………………………………

168 -62.9 42.7 261.1 29.1 6.36 -94.7	38.7	26.0	271.4	1661.3	185.4

192 -62.9 42.7 261.1 29.1 6.36 -94.7	38.7	26.0	271.4	1661.2	185.3

216 -62.9 42.7 261.1 29.1 6.36 -94.7	38.7	26.0	271.4	1661.1	185.3

240 -62.9 42.7 261.1 29.1 6.36 -94.7	38.7	26.0	271.4	1661.1	185.3

*	na0	k0	cl0	B0	kv	na	k	cl	beta gamma

*	140.0	5.8 116.0 228.2 1.580	60.0 232.3	71.1 0.039 1.50

*	pna	pk	pcl	inc	ikc	inkcc	hp	kb

*	0.00170 0.01150 0.01100 0.00007 0.0000800 0.0000000080	240 0.0

Net_flux	PUMP	Channel	NC	KC	NKCC

*	Na -1.6635 0.6018 1.0499 0.0000 0.0119

*	K	1.1090 -0.5666 0.0000 -0.5543 0.0119

*	Cl	0.0000 -0.5193 1.0499 -0.5543 0.0238

*	Influx	PUMP	IChannel INC	IKC	INKCC

*	Na	0.0000 0.6197 1.1368 0.0000 0.0874

*	K	1.1090 0.1737 0.0000 0.0538 0.0874

*	Cl	0.0000 0.3146 1.1368 0.0538 0.1748

*	Efflux	PUMP	EChannel ENC	EKC	ENKCC

*	Na -1.6635 -0.0179 -0.0869 0.0000 -0.0755

*	K	0.0000 -0.7403 0.0000 -0.6082 -0.0755

*	Cl	0.0000 -0.8339 -0.0869 -0.6082 -0.1511

z	OSOR (A/V)*1000

*	-1.75 3.52 157.16
